# Multiscale interactions between plant part and a steep environmental gradient determine plant microbial composition in a tropical watershed

**DOI:** 10.1101/2020.07.20.212811

**Authors:** Jared Bernard, Christopher B. Wall, Maria S. Costantini, Randi L. Rollins, Melissa L. Atkins, Feresa P. Cabrera, Nicolas D. Cetraro, Christian K. J. Feliciano, Austin L. Greene, Philip K. Kitamura, Alejandro Olmedo-Velarde, Vithanage N. S. Sirimalwatta, Helen W. Sung, Leah P. M. Thompson, Huong T. Vu, Chad J. Wilhite, Anthony S. Amend

**Affiliations:** Department of Plant and Environmental Protection Sciences, University of Hawai‘i–Mānoa, Honolulu, Hawai‘i 96822, USA; Hawai‘i Institute of Marine Biology, University of Hawai‘i– Mānoa, Kāneʻohe, Hawai‘i 96744, USA; Pacific Biosciences Research Center, University of Hawai‘i–Mānoa, Honolulu, Hawai‘i 96822, USA; Department of Biology, University of Hawai‘i–Mānoa, Honolulu, Hawai‘i 96822, USA; Department of Botany, University of Hawai‘i–Mānoa, Honolulu, Hawai‘i 96822, USA; Department of Natural Resources and Environmental Management, University of Hawai‘i–Mānoa, Honolulu, Hawai‘i 96822, USA; Department of Molecular Biosciences and Bioengineering, University of Hawai‘i–Mānoa, Honolulu, Hawai‘i 96822, USA

## Abstract

Plant microbiomes are shaped by forces working at different spatial scales. Environmental factors determine a pool of potential symbionts while host physiochemical factors influence how those microbes associate with distinct plant tissues. Interactions between these scales, however, are seldom considered. Here we analyze epiphytic microbes from nine *Hibiscus tiliaceus* trees across a steep environmental gradient within a single Hawaiian watershed. At each location we sampled eight microhabitats: leaves, petioles, axils, stems, roots, and litter from the plant, as well as surrounding air and soil. While the composition of microbial communities is driven primarily by microhabitat, this variable predicted more than twice the compositional variance for bacteria compared to fungi. Fungal community compositional dissimilarity increased more rapidly along the gradient than did bacteria. Additionally, the spatial dynamics of fungal communities differed among plant parts, and these differences influenced the distribution patterns and range size of individual taxa. Within plants, microbes were compositionally nested such that aboveground communities contained a subset of the diversity found belowground. Our findings identify potential differences underlying the mechanisms shaping communities of fungi and bacteria associated with plants, and indicate an interaction between assembly mechanisms working simultaneously on different spatial scales.

## Introduction

Plants harbor communities of microorganisms that influence their biology, including phenology [1], water conductance [2], niche occupancy and range expansion [3–5], and competitive ability [6]. Nearly all plant traits are likely affected by microbial partners. Our understanding of symbiotic microbial functions, however, has outpaced our understanding of how plants and their microbes form relationships that persist in nature. This disjuncture stems from the sheer complexity of microbial communities, and is compounded by assembly patterns governed by interacting processes at multiple ecological scales, from the landscape level to variation among plant tissues within an individual plant [7, 8].

At large scales, abiotic environmental factors influence plant microbiome composition [9]. Plant-associated microbial communities can change across elevation gradients [10–14] or in response to soil properties (*e.g.*, organic carbon, soil pH, nitrogen content [15] and land-use history [16]). Even in the absence of obvious environmental clines, geographic distance coupled with presumed dispersal limitation can alter microbiome composition [17–19]. Host identity [20] and genotype [8, 21, 22] can also covary with environmental factors, which can influence microbial composition as well [23].

Within plants, distinct microbial communities associate with different plant tissues [24–26] and this factor can be more predictive of plant microbial composition than location, even over broad geographic areas. For instance, a root microbiome can be more similar to another root microbiome several hundred kilometers away than to a leaf microbiome on the same individual plant [23]. The assembly processes governing within-plant microbial assembly is not merely a recapitulation of dispersal- or environmentally-mediated dissimilarity at a smaller scale. While large environmental clines can result in compositional turnover (*i.e.*, replacement of species between communities) [27], microbiomes within an individual plant tend to be compositionally nested such that apical parts (*e.g.*, leaves, flowers) house a subset of the microbial species associated with subterranean plant parts, which themselves are most species rich [19]. This nestedness pattern indicates that, although compositionally distinct, microbiomes of different plant tissues within a plant are neither entirely independent of each other nor of the larger environmental species pool. Therefore, the microbial composition of a given plant part may be partially attributed to the local abiotic environment [28] as well as to strong selective forces from the host plant itself [29].

A multiscale model of plant microbiome assembly might include the interactions of abiotic environmental factors circumscribing a local pool of microbial symbionts as a higher-order term [30–32], which is secondarily partitioned *within* plants, via a distinct set of processes that shape plant microbiomes across host tissues [24–26, 29, 33]. Despite a large and growing body of literature parameterizing plant microbial assembly at landscape and within-plant scales [23, 28, 33–35], few have considered how large-scale assembly patterns impact within-plant microbiome composition, or how residency within plant microhabitats relates to microbial distributions at landscape scales. In other words, to what extent do these scale-relevant processes interact?

Here we sampled epiphytic fungi and bacteria on multiple tissues of *Hibiscus tiliaceus* trees as well as adjacent soil, leaf litter, and air microhabitats across a steep environmental gradient on the island of Oʻahu, Hawaiʻi. We hypothesized that plant tissue type would determine microbial composition within individual plants, and that microbial compositional differences across the gradient would differ among microbiomes from different plant tissues due to differential specificity and dispersal ability of community members [36]. Specifically, we hypothesized that host-independent microbial communities associated with air and soil would demonstrate comparatively less distance decay (*i.e.*, compositional dissimilarity over distance) because their distribution ought not to be buffered by specificity to hosts. Second, we assess the interaction between small and large scales by testing the hypothesis that microbes with broader occupancy among plant microhabitats will have a greater range along the gradient. Next, we assess whether compositional nestedness within plants varies with the abiotic environment. Last, we evaluate whether fungi and bacteria differ fundamentally in their distribution and assembly across plant parts due to differences in dispersal ability and microhabitat specificity.

## Materials and Methods

### Site description and sampling

We collected microbial samples from, and adjacent to, nine mature healthy *Hibiscus tiliaceus* L. (Malvaceae; Hawaiian: *hau*) individuals within Waimea Valley on the north shore of the island of Oʻahu, Hawaiʻi (Fig. 1). *Hibiscus tiliaceus* is a medium sized tree or shrub with broad heart-shaped leaves. It is indigenous to the tropical Indo-Pacific but has attained a pantropical distribution [37], and was possibly introduced to Hawaiʻi by Polynesian settlers [38]. Sample trees were located along a transect that paralleled a perennial stream. Locations of *H. tiliaceus* ranged from 0–700 m elevation, spanning 1174–2026 mm annual rainfall over a distance of ~5.5 km (Fig. 1). Rainfall, elevation and distance from the shore are highly collinear so we refer to this continuous variable cline as the “environmental gradient”. For each plant, we sampled epiphytic microbes from standardized surface areas (200 cm^2^) of leaves, petioles, axils, stems, roots and adhering soil, and litter with a sterile cotton swab (one per microhabitat). We sampled surface microbes because 1) doing so let us standardize sampling area across these microhabitat types; 2) it mitigates known issues of co-amplification of chloroplasts (16S) and plant host (ITS) DNA when microbial:plant biomass ratios are low; and 3) plant surfaces represent important interfaces between plants and their environments where gas, water and nutrient exchange occurs.

**Fig. 1.**
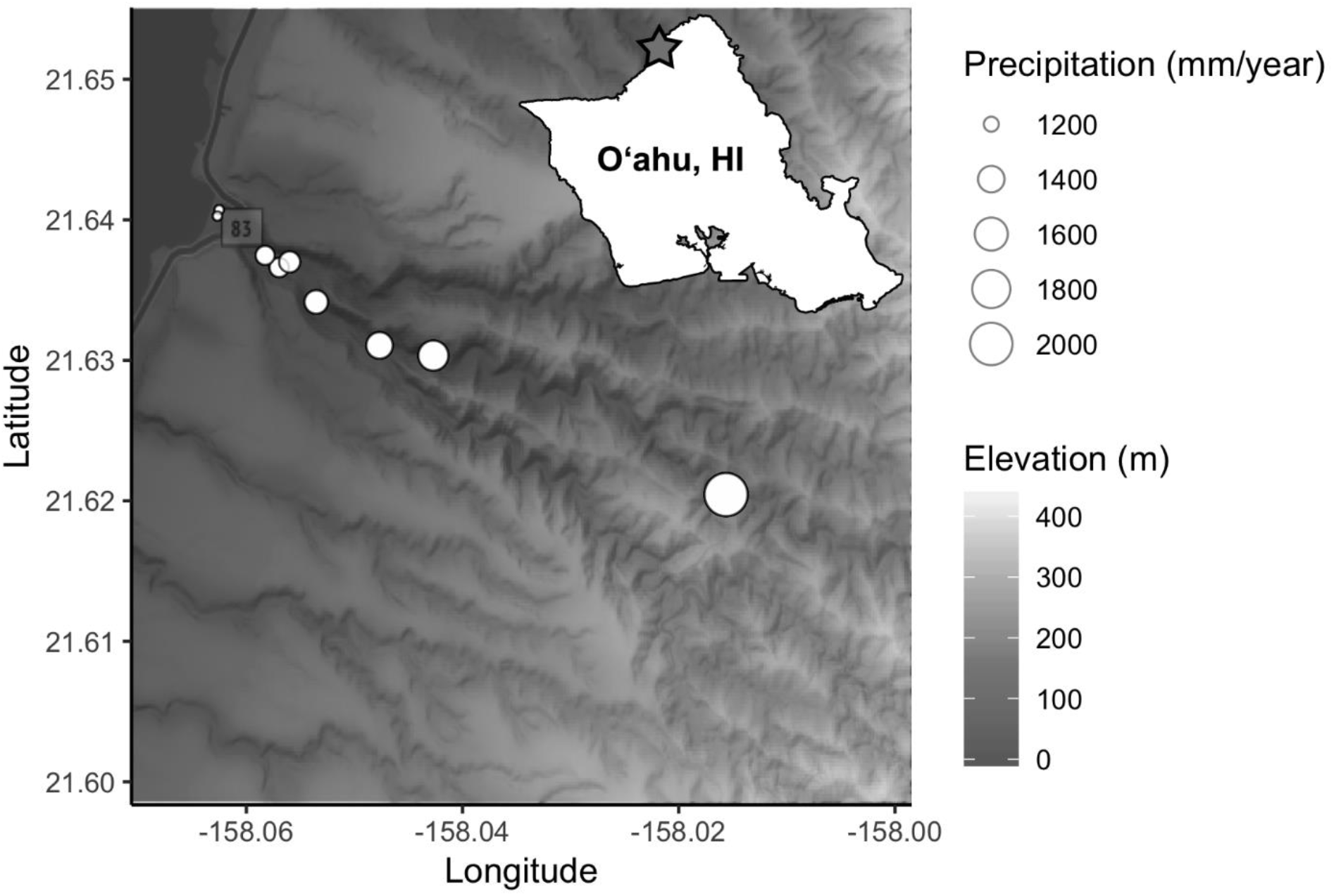
Locations of sampled *Hibiscus tiliaceus* in Waimea Valley, Oʻahu (inset), denoted by white circles. Circle radius indicates the amount of annual precipitation, with shaded gradient indicating elevation. Both increase as a function of distance from shore.

To sample the soil microbiome, we dug a 5-cm hole using a sterilized corer ~1 m from the canopy edge to avoid sampling the rhizosphere, and swabbed the sides and bottom of the hole. For two weeks prior to collecting the above samples, we also deployed an air sampler modified from Quesada *et al.* [39] at each of our *H. tiliaceus* sites, which collected aerial microbes on rotating sterilized glass slides lined with microtiter plate sealing film (Thermo Fisher Scientific, Waltham, MA). We immediately transferred all swabs and film to a microcentrifuge tube with 1 mL lysis buffer and garnet homogenization beads (Qiagen NV, Venlo, Netherlands; see Supplementary Methods for details). In total, we collected 72 biological samples (9 sites by 8 microhabitats). For negative controls, we exposed sterile swabs dipped in lysis buffer to the air for ~20 sec. and processed them in the same way as the biological samples. A single air sampling film that we did not expose to the environment served as an air sampler negative control. Hereafter, we refer to the 5 plant tissue types (leaves, petioles, axils, stems, roots) as well as the air, litter, and soil collectively as “microhabitats,” which we treat as categorical variables.

### DNA extraction and library preparation

We extracted and purified DNA using the KingFisher™ Flex™ System (Thermo Fisher Scientific, Waltham, MA, USA) with a MagAttract PowerSoil KF Kit (Qiagen NV, Venlo, Netherlands) following the manufacturer’s protocol, but increased the volume of elution buffer to 125 μL. To survey bacteria, we amplified the V4 region of the 16S ribosomal RNA gene with the primers 515F and 806R, and for fungi we amplified the ribosomal internal transcribed spacer 1 (ITS1) using the forward ITS1F and reverse ITS2 primer schemes [40, 41]. Both library preparations involved an 8-base-pair index, a gene-specific primer, and an adaptor construct concatenated on the same oligo. PCR protocols entailed an initial phase of 95° C for 3 min using Hot Start *Taq* polymerase (New England BioLabs Inc., Ipswich, MA, USA), followed by 35 cycles at 50°C and 52°C annealing temperatures for 16S and ITS amplicons respectively. We then cleaned and normalized PCR products using the Just-a-Plate™ 96 PCR Purification and Normalization Kit (Charm Biotech, San Diego, CA, USA). We separately pooled bacteria and fungi amplicons and checked the quality of libraries with a Bioanalyzer 2100 (Agilent Technologies, Santa Clara, CA, USA). The Advanced Studies in Genomics, Proteomics and Bioinformatics Laboratory at the University of Hawai‘i – Mānoa sequenced the libraries in two runs using separate 300-bp paired-end Illumina MiSeq v3 reagent kits (Illumina Inc., San Diego, CA, USA). Sequence data are available at the Sequence Read Archive as PRJNA543421.

### Data processing and analyses

Because of fundamental differences between ITS and 16S loci, we used different processing pipelines. Due to heterogeneity in ITS amplicon lengths, we did not attempt to pair ITS reads. We processed read 1 fungal FASTQ files using ITSxpress [42] to isolate ITS1 regions of variable lengths from adjacent conserved ribosomal subunit genes. We used the FASTX-Toolkit [43] to filter sequences by quality scores, and to discard reads that met at least one of the following conditions: 1) 10% or more of their bases contained a *q*-score lower than 25; 2) they contained an “N” nucleotide; or 3) their length was less than 20 bp. We used VSEARCH [44] to identify and remove chimeras. The DADA2 package [45] corrected sequencing errors and binned amplicon sequence variants (ASVs; see Callahan *et al.* [46] for discussion on ASVs). We used CONSTAX [47] with the UNITE 8.0 database to assign fungal taxonomy via consensus of three separate classification algorithms. Finally we used the LULU algorithm [48] to collapse putative within-genome ribotype variants into a single ASV.

We used DADA2 to process 16S reads. First we truncated reads at position 210 (190 for the reverse read) and discarded these if they contained at least one base below quality 2 or a number of expected errors above 3. We used DADA2’s default parameters to denoise the data, and merged reads if they overlapped by at least 20 bases, allowing for one mismatch at most. We used MOTHUR [49] along with the Silva database v132 to filter and annotate sequences. We removed potential chimeras with VSEARCH as implemented in MOTHUR, and assigned bacteria taxonomy via the MOTHUR functions classify.seqs() and classify.otus(). As with the ITS pipeline, we corrected bacteria ASVs with LULU. Finally, we used the R package decontam [50] to identify and remove putative contaminants (33 bacteria and 4 fungi) based on their prevalence in extraction and PCR negative controls.

The processed dataset contained 15 649 bacterial and 12 558 fungal ASVs, belonging to 96 and 35 classes respectively. We amalgamated ASV read abundance data, taxonomic assignments and sample metadata using the R package phyloseq [51], and appended tree location climate information from University of Hawai‘i–Mānoa’s Climate of Hawaiʻi data portal [52] using the R package raster [53].

To test the extent to which either location along the gradient or microhabitat explained the composition of bacteria or fungi, we used permutational multivariate analysis of variance (PERMANOVA) [54] on a Bray-Curtis dissimilarity matrix of Hellinger transformed data with 10,000 permutations in the R package vegan [55]. We used the same distance matrices in a series of Mantel tests examining the correlation between community dissimilarity and geographic distance (*i.e.*, pairwise distance between tree locations) for each microhabitat separately and combined, using Bonferroni-corrected *p*-values to account for Type 1 errors due to multiple hypothesis testing [56]. To examine the extent to which responses of communities of fungi and bacteria were coordinated in response to within- and among-plant factors, we tested the correlation between distance matrices for fungi and bacteria samples using Mantel.

For each site and all sites combined, we used the R packages vegan and bipartite [57, 58] to evaluate nestedness in two ways: 1) a temperature statistic (*T*) in which 0° is perfectly nested and 100° is perfectly random based on pairwise compositional differences [59]; and 2) the Nestedness metric based on Overlap and Decreasing Fill (NODF) [60]. Furthermore, we assessed relationships between nestedness metrics and gradient location (distance from shore) using Spearman correlations. To account for differences in sampling depth, these analyses used sample counts randomly down-sampled (rarefied) to the same depth per sample (bacteria: 26 406, fungi: 20 000).

To examine the relationships between local ASV abundances, geographic ranges and number of occupied microhabitats, we used regression analyses. We calculated local abundance as the mean sequence abundance per sample of an ASV where present, omitting nulls. The maximum distance between locations where we detected an ASV along the transect gave us its range; ASVs detected at a single location had a range of zero. We determined an ASV’s habitat occupancy as the sum of microhabitats where we detected an ASV. Although these calculations may include dead cells and thus overestimate empirical ranges or habitat breadths of viable cells, we believe these errors will be propagated randomly throughout the dataset and are nevertheless preferable to arbitrary sequence abundance cutoffs. A two-tailed *t*-test assessed the difference of mean range sizes between distributions of fungi and bacteria. We furthermore calculated the range distribution of microbial genera to compare 16S and ITS marker genes at comparable taxonomic levels. Scripts, data and markdown documents necessary to reproduce all analyses are accessible on GitHub (github.com/cbwall/Waimea-plant-microbiomes).

## Results

PERMANOVA tests indicate that communities were significantly differentiated by microhabitat type, with more variance explained for bacteria than fungi (Table 1). Composition of bacteria and fungi were also explained by location, but to a lesser extent. At the class level, communities of bacteria from above- and belowground samples were compositionally distinct (Supplementary Fig. S1) whereas fungal classes were uniformly distributed among microhabitats (Supplementary Fig. S2). At the ASV scale, however, compositional differences were more readily apparent among bacteria samples from different microhabitats and between fungal samples from below- and aboveground (Fig. 2).

**Table 1.**
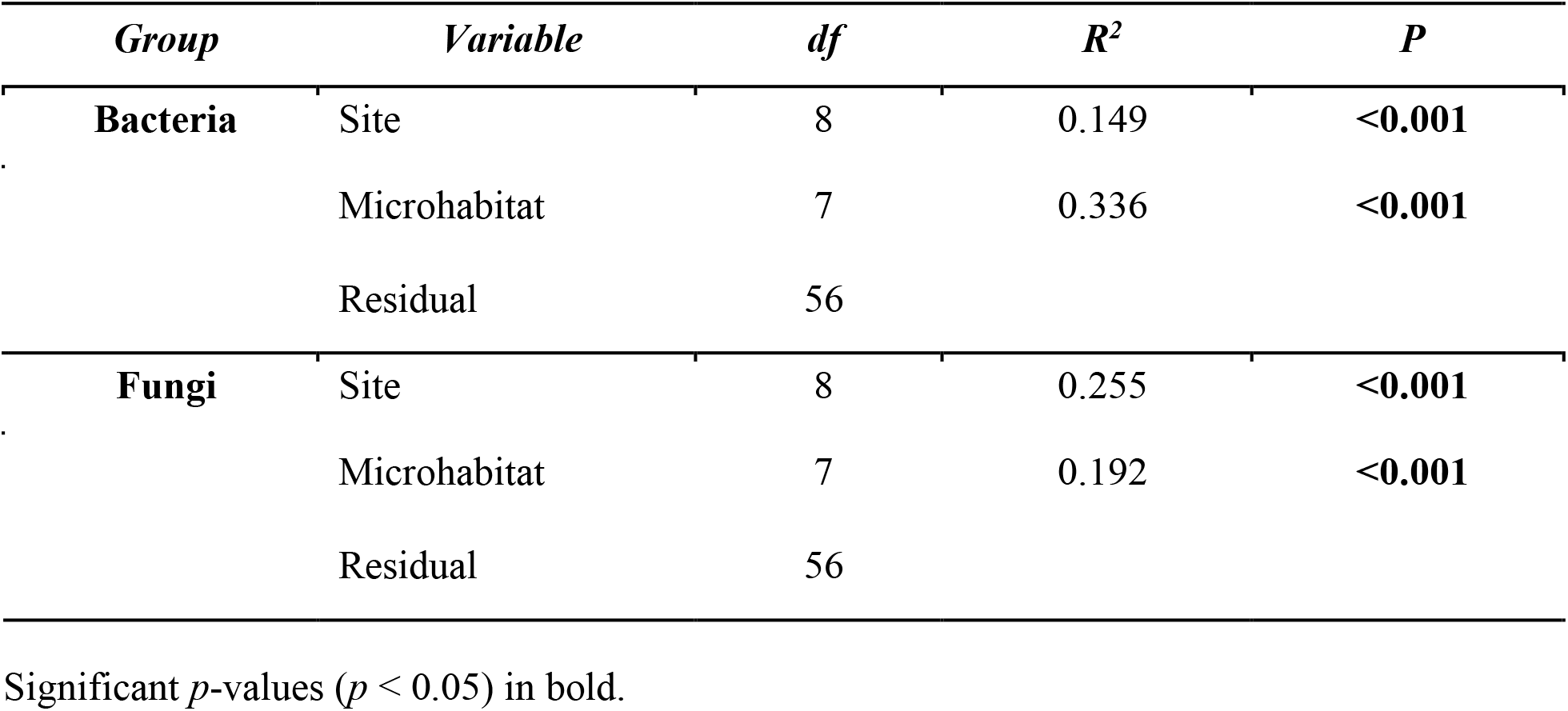
PERMANOVA results showing the ability of variables to explain compositional variance.

**Fig. 2.**
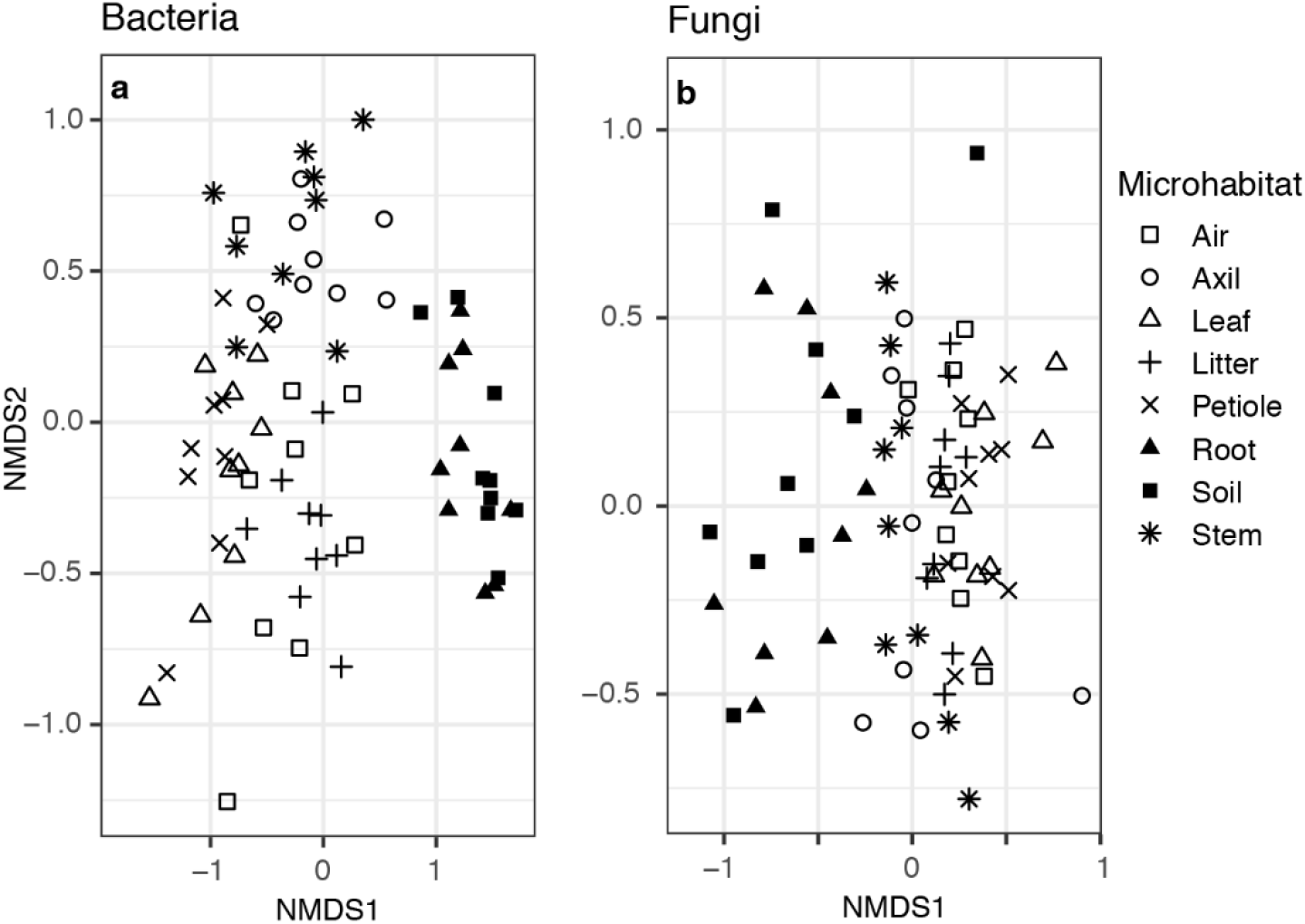
Non-metric multidimensional scaling plots of microbial community dissimilarity with shapes indicating microhabitat type. (a) Bacterial communities cluster by above- and belowground microhabitats. (b) Fungal communities also cluster somewhat by aboveground and belowground parts, albeit with higher dispersion in the belowground communities and more overlap with the aboveground parts compared to communities of bacteria.

There was no evidence for a distance decay relationship among bacterial communities, except for those associated with roots (Table 2). Conversely, communities of fungi, when aggregated by either site or microhabitat, demonstrated significant distance decay patterns over the environmental gradient. Significance and variance explained by the Mantel test differed among fungal communities sampled from different microhabitats, indicating an interaction between microhabitat and environment. Mantel correlations of subterranean fungal samples (roots and soil) showed the steepest slope, while results for fungal communities on axils and leaves were not significant (Table 2).

**Table 2.**
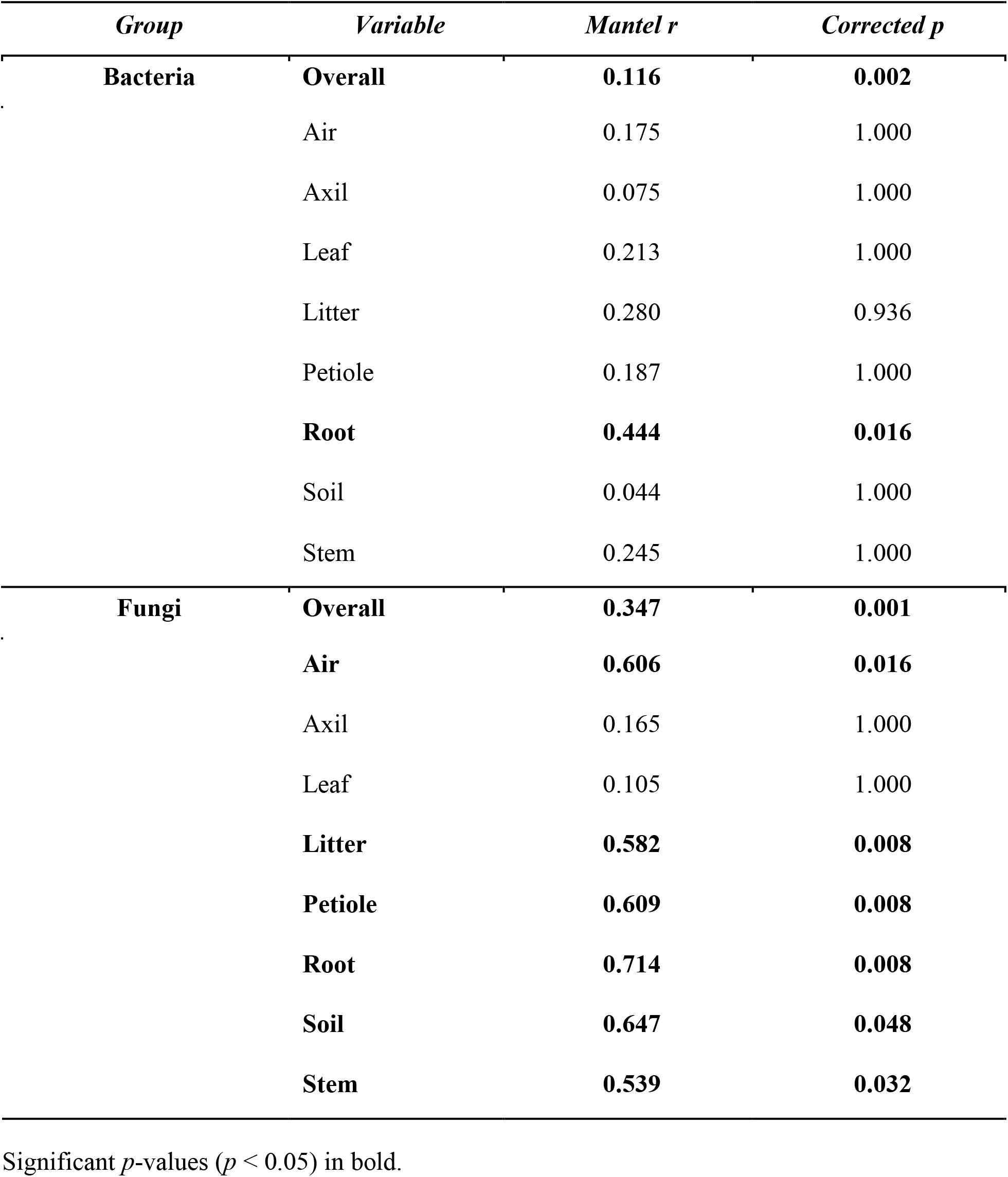
Mantel test of relationships between pairwise community composition dissimilarity and geographic distance between communities, both overall and per microhabitat.

Despite differences between the relative importance of drivers of fungi and bacteria, compositional dissimilarity between samples was correlated. In other words, fungal and bacterial community composition, as assessed by Bray-Curtis dissimilarity, was significantly more correlated than would be expected by chance (Mantel *r* = 0.417, *p* = 0.001; Supplementary Fig. S3).

Communities of both bacteria and fungi were significantly nested by microhabitat (Fig. 3), with belowground samples containing much of the species diversity found within the other microhabitat communities. Each had a moderate nestedness temperature (bacteria: *T* = 38.8°, *p* = 0.001, NODF = 39.039%; fungi: *T* = 43.1°, *p* = 0.001, NODF = 34.760%). We detected no directional nestedness with regard to location along the gradient (*e.g.*, microbial communities in lower elevation locations were not subsets of those in higher locations).

**Fig. 3.**
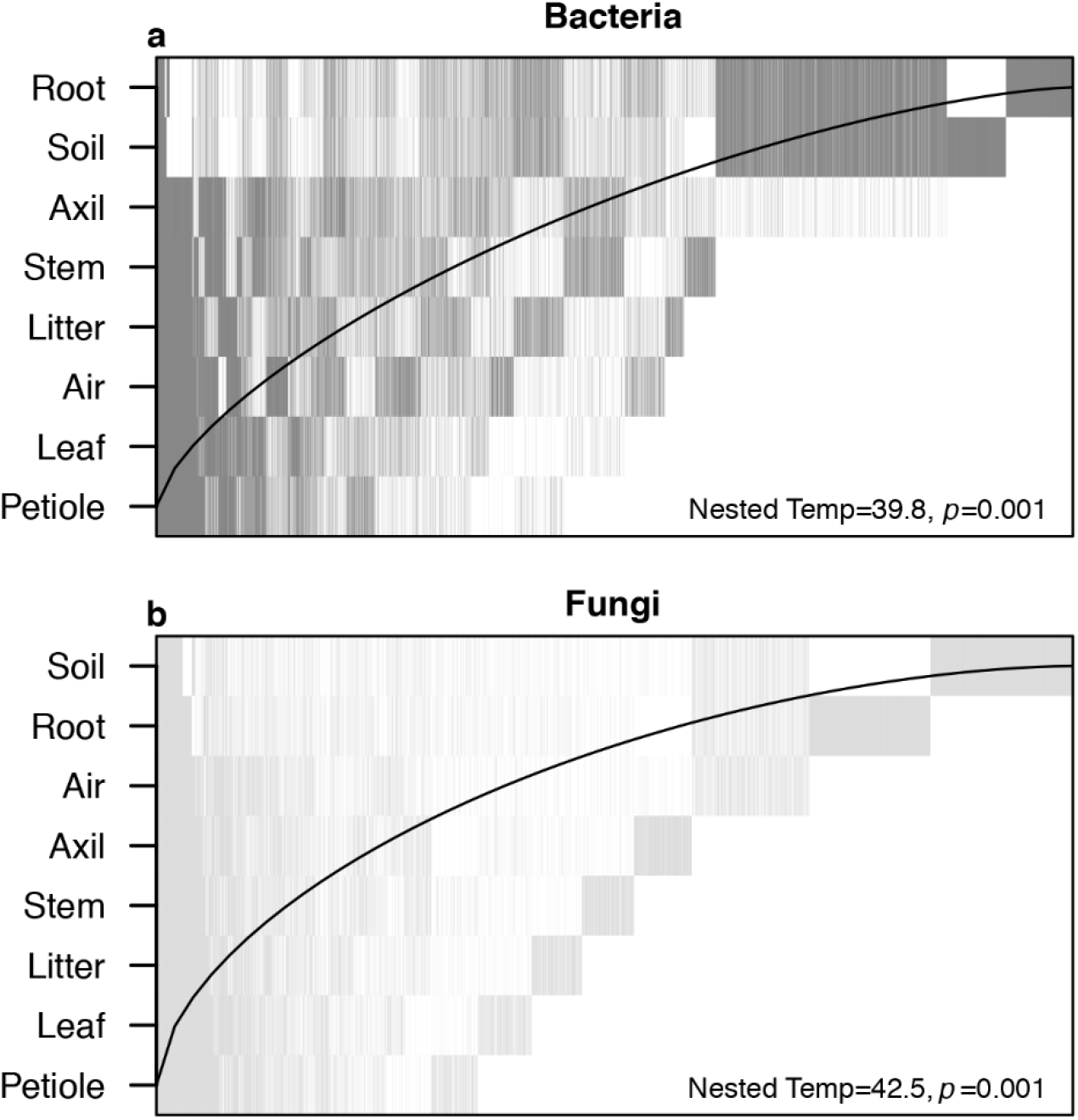
Nestedness plots of overall (a) bacterial and (b) fungal composition aggregated by microhabitat type, where each vertical line represents an ASV’s presence among microhabitats. Both fungi and bacteria display moderate nestedness with most of the microbial diversity contained in the subterranean microhabitats. In a perfectly nested model (*T* = 0°) ASVs would be constrained to the left of the black line.

The range size of bacterial and fungal ASVs differed significantly (*t* = 32.062, df = 13235.00, *p* < 0.001), with the mean range of bacteria (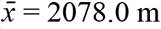, SD = 2044.9 m) being nearly twice the range of fungi (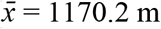, SD = 1845.1 m; Fig. 4). As expected, the range sizes of microbial genera were larger than those of ASVs, but bacteria (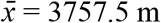, SD = 2096.5 m) remained significantly more widespread than fungi (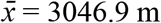, SD = 2270.3 m; *t* = 5.729, df = 869.79, *p* < 0.001). For both bacteria and fungi, an ASV’s range size correlated positively with the number of microhabitats in which an ASV occurred, although this relationship was stronger for fungi (*R*^*2*^ = 0.451, *p* < 0.001) compared to bacteria (*R*^*2*^ = 0.262, *p* < 0.001). Fungal ASV abundance scarcely correlated with range size (*R*^*2*^ = 0.012, *p* < 0.001), and not at all for bacteria (*p* = 0.554).

**Fig. 4.**
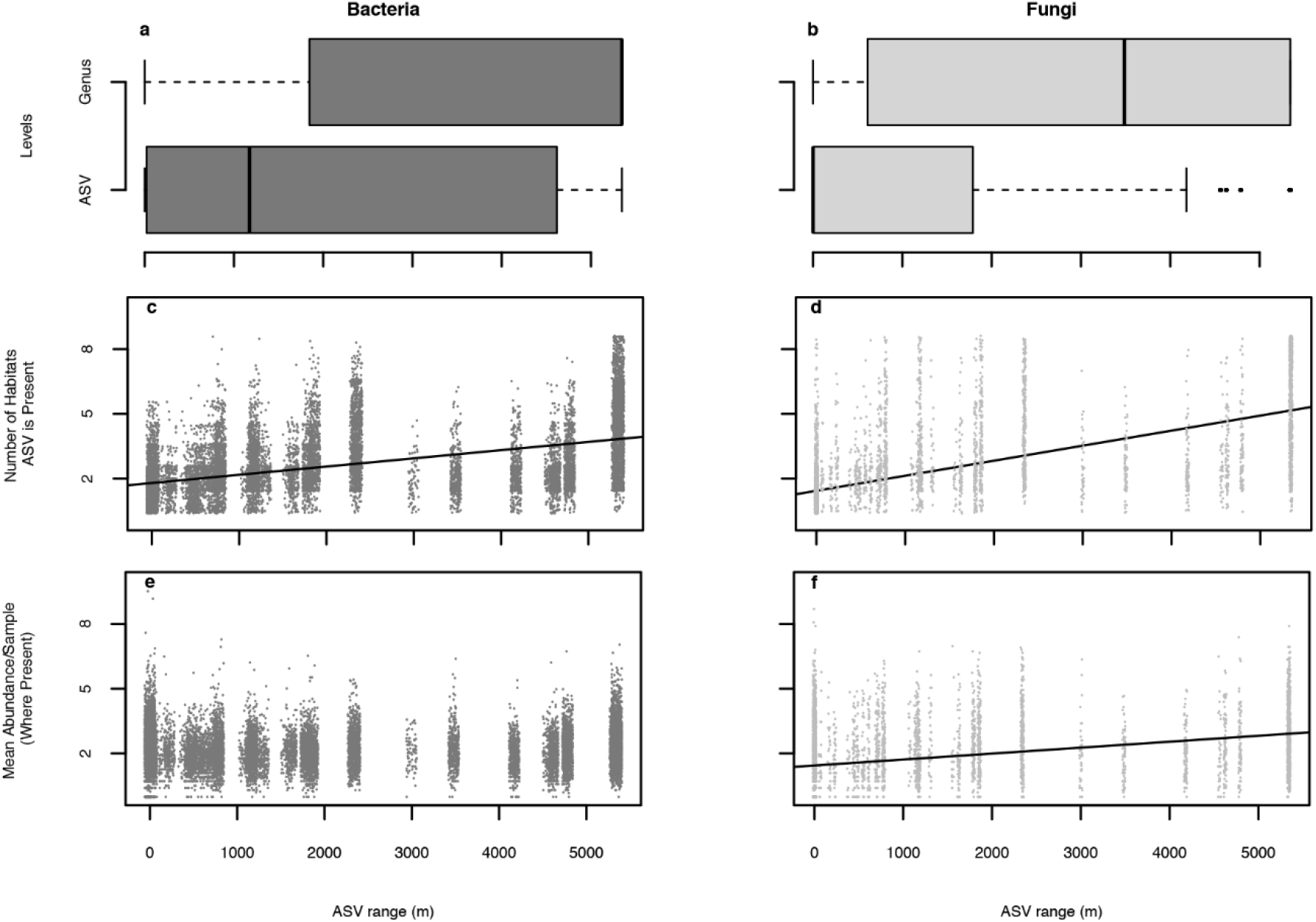
Range spans of (a) bacteria and (b) fungi genera and ASVs across Waimea Valley, Oʻahu. Boxes contain the interquartile range, the line depicts the median and the whiskers contain 95^th^ percentiles. Lines depict the Pearson`s least-squares fit of the relationship between the number of microhabitats in which each (c) bacterial and (d) fungal ASV occurs and its range within the study. The non-zero mean abundance of each ASV per sample is not significantly predicted by that ASVs range in bacteria (e) but is weakly related to ASV range in fungi (f), where abundances are log-transformed.

## Discussion

Here we demonstrate that small and large scale factors shape the plant microbiome in distinct and synergistic ways. Fungal communities responded more to location along the environmental gradient (large scale variable) than to microhabitat type (small scale variable). Nonetheless, strong interactions between these variables influenced fungal compositions. Specifically, fungal communities had a high variation of distance decay patterns among microhabitat types. Bacteria, in contrast, were widely distributed along the gradient, but also sharply partitioned by microhabitat. Moreover, microhabitat generality was a significant predictor of range sizes of microbes, although nearly twice as predictive for fungal ASVs as for bacterial ones. Despite these seemingly divergent compositional patterns, dissimilarity between biological samples was significantly correlated between fungi and bacteria, and both were compositionally nested within plants, suggesting that at least some drivers of community assembly apply universally to members of both kingdoms.

### Effect of microhabitat on microbial distribution

Similar to previous studies [19, 28, 61], we show that microhabitat type significantly influences microbial community composition. This factor is twice as strong for bacteria as fungi (Table 1). At the ASV level, both bacterial and fungal communities are distinguished primarily by whether they are located above- or belowground (Fig. 2). While this distinction is still apparent at the class level for bacteria (Supplementary Fig. S1), nearly all fungal classes were uniformly distributed across the eight microhabitats (Supplementary Fig. S2).

Our study focused on epiphytic microbiomes, which may have important implications for determining patterns of microhabitat niche breadth. Previous studies showed that different variables drive epiphytic versus endophytic microbes [23]. Because surface communities contain a portion of microbes only transiently associated with hosts, more persistent endophytic microbes might demonstrate higher fidelity than would epiphytes. In a survey of endophytic fungi associated with grassland plants, Wearn *et al.* [29] found most taxa were associated with a single plant tissue, and that those associated with more than one were rare, implying a link between specialization and endophytic microhabitat [33].

The wide niche breadth of some microbes is reflected in patterns of compositional nesting, with aboveground communities comprising a subset of the ASVs present belowground (Fig. 3). This pattern is replicated at all locations throughout our environmental gradient, and is consistent with previous studies on other plant species [19, 28, 61]. Although the causes of these vertically stratified patterns are undetermined, soils may inoculate plants with generalist microbes during germination, followed by additional non-random factors (*e.g.*, desiccation, dispersal limitation) that subset the original microbial community over time and distance from the ground [19, 61]. Intriguingly, rhizosphere communities, not soil, are basal in the nestedness hierarchy, congruent with previous studies [19, 27]. This might reflect this microhabitat’s unique mix of diverse soil- and plant-associated microbial communities in a resource rich and heterogeneous environment.

### Interactions between microhabitat and environmental gradient

In contrast to communities of bacteria, fungi demonstrated a strong interaction between microhabitat type and compositional dissimilarity along the environmental gradient (Table 2). Our results show that subterranean fungal communities had the steepest dissimilarity slopes, which could result from greater heterogeneity among subsurface microhabitats [61–63] or more limited dispersal among soil-bound microbes [36] as compared to aboveground counterparts. The absence of distance decay patterns among foliar microbiomes is not surprising given the high compositional variance of these consortia compared with those on other plant surfaces [19, 64] even within plant individuals [26]. Although we originally suspected that communities on plant tissues might demonstrate less compositional turnover than free-living samples [27], and that leaf litter communities might be intermediate, we did not find evidence for such a pattern. Instead fungal communities in air, which we suspected should be the least constrained by either dispersal or environment, showed a strong distance decay pattern (Table 2).

Furthermore, the correlation between an ASV’s range size and the number of microhabitats it occupies (Fig. 4) implies a relationship between microhabitat specialization and environmental (in)tolerance, such that niche breadth within a plant is predictive of ecological breadth across the entire gradient. Some environmental gradients across Waimea Valley might be replicated at a small scale within a plant (moisture being the most obvious), so this relationship might reflect environmental filtering for certain traits (*e.g.*, desiccation tolerance). In fact, phylogeny and physiology can dictate microbial biogeography on a global scale [65]. Alternatively, taxa with broad niches have a higher probability of establishing after dispersal as they are more likely to find a suitable microhabitat. In our study, the most widespread bacterium (Rokubacteria) and the most widespread fungus (Geminibasidiomycetes) are known as microhabitat generalists, with the latter being both heat and desiccation tolerant (Supplementary Figs S4 & S5; [66, 67]).

The weak (fungi) or non-significant (bacteria) relationship between ASV abundance and microhabitat occupancy suggests that the correlations with niche breadth at small and large scales are not solely attributable to numerical dominance and ascertainment biases. Despite having wider ranges, generalists are not, on average, more abundant than less generalist taxa, suggesting that neither generalists nor more specialized taxa dominate the communities in our study system (Fig. 4).

### Differences and similarities between bacteria and fungi

Several factors might explain why the environmental gradient impacted communities of fungi more strongly than bacteria. Dispersal limitations [9, 68], environmental dissimilarity [69], host genetic clines [8, 21] or some combination of these or other factors may affect bacteria and fungi differently. Of the relatively few studies that concurrently examine bacteria and fungi, fungal communities tend to be more influenced by geographic distance. Coleman-Derr *et al.* [23] found that agave hosts in southern California and Mexico shared a smaller percentage of fungi compared to bacteria (18.2% and 72.2% respectively). A study on *Populus deltoides* roots in Tennessee and North Carolina revealed that geography explained more variance for epiphytic fungal communities than for bacterial ones, although this trend was not the same for endophytes [70].

Notably, differing rates of evolution between marker genes might partially explain the contrasting patterns we found between fungi and bacteria. Because the ITS gene is less phylogenetically conserved than the adjacent small subunit rRNA gene, it circumscribes taxa at a finer scale than does 16S. ASVs derived from 16S and ITS sequences, therefore, may not be equivalent in terms of evolutionary divergence.

Notwithstanding differences in spatial patterns and plant part specialization, however, the community compositional turnover of fungi was correlated with that of bacteria (Supplemental Fig. S3). Regardless of whether this relationship reflects interkingdom microbial biotic interactions, or merely coordinated response to environmental cues, we find it remarkable that despite billions of years of evolutionary divergence, organisms’ distributions are similarly constrained by being diminutive and plant-associated.

## Conclusion

Different factors govern the distribution of tree-associated bacteria and fungi along a steep environmental gradient. Microhabitat type (*i.e.*, soil, litter, plant parts, air) was a strong driver of bacterial communities, whereas communities of fungi were more influenced by location along the environmental gradient, and its interaction with microhabitat. These findings suggest differences in the underlying mechanisms shaping fungal or bacterial communities associated with plants, which could be based on a combination of dispersal abilities, and environmental or microhabitat specificity. Despite these differences, communities from both microbial domains were strongly nested such that aboveground microbes represented a subset of subterranean ones. Together these results indicate that microbial assembly on plants works simultaneously on multiple interacting scales, a complex dynamic that should be accounted for, explicitly when determining the drivers of plant symbionts.

## Acknowledgments

We greatly appreciate Josie Hoh and Chad Durkin at the Waimea Botanical Garden, Kirsten Cannoles and Sean Swift for their assistance in the laboratory, and Cédric Arisdakessian and Mahdi Belcaid for developing the bioinformatic pipeline used. We also acknowledge the generous support of Illumina Corporation, Michele Langner, and the ASGPB for sequencing. This work was supported by a grant from the W.M. Keck foundation to A.S.A.

## Author Contributions

A.S.A planned and designed research, and organized funding; all authors conducted fieldwork, analyzed data, and contributed to manuscript content; V.N.S.S. prepared libraries; C.B.W. and J.B. curated codes; A.S.A., J.B. and C.B.W. interpreted results; and J.B., C.B.W., M.S.C., R.L.R. and A.S.A. wrote the manuscript.

## References

1. Zahn G, Amend AS. Foliar fungi alter reproductive timing and allocation in *Arabidopsis* under normal and water-stressed conditions. Fungal Ecol 2019; 41: 101–106.

2. Arnold AE, Engelbrecht BMJ. Fungal endophytes nearly double minimum leaf conductance in seedlings of a neotropical tree species. J Trop Ecol 2007; 23: 369–372.

3. Taghavi S, Garafola C, Monchy S, Newman L, Hoffman A, Weyens N, et al. Genome survey and characterization of endophytic bacteria exhibiting a beneficial effect on growth and development of poplar trees. Appl Environ Microbiol 2009; 75: 748–757.

4. Berg G, Köberl M, Rybakova D, Müller H, Grosch R, Smalla K. Plant microbial diversity is suggested as the key to future biocontrol and health trends. FEMS Microbiol Ecol 2017; 93.

5. Choudoir MJ, Barberan A, Menninger HL, Dunn RR, Fierer N. Variation in range size and dispersal capabilities of microbial taxa. Ecology 2017; 99: 322–334.

6. Klironomos JN. Feedback with soil biota contributes to plant rarity and invasiveness in communities. Nature 2002; 417: 67–70.

7. Dini-Andreote F, Raaijmakers JM. Embracing community ecology in plant microbiome research. Trends Plant Sci 2018; 23: 467–469.

8. Beilsmith K, Thoen MPM, Brachi B, Gloss AD, Khan MH, Bergelson J. Genome-wide association studies on the phyllosphere microbiome: embracing complexity in host-microbe interactions. Plant J Cell Mol Biol 2019; 97: 164–181.

9. Peay KG, Kennedy PG, Talbot JM. Dimensions of biodiversity in the Earth mycobiome. Nat Rev Microbiol 2016; 14: 434–447.

10. Wang J, Soininen J, He J, Shen J. Phylogenetic clustering increases with elevation for microbes. Environ Microbiol Rep 2012; 4: 217–226.

11. Zimmerman NB, Vitousek PM. Fungal endophyte communities reflect environmental structuring across a Hawaiian landscape. Proc Natl Acad Sci 2012; 109: 13022–13027.

12. Yang Y, Gao Y, Wang S, Xu D, Yu H, Wu L, et al. The microbial gene diversity along an elevation gradient of the Tibetan grassland. ISME J 2014; 8: 430–440.

13. Shen C, Ni Y, Liang W, Wang J, Chu H. Distinct soil bacterial communities along a small-scale elevational gradient in alpine tundra. Front Microbiol 2015; 6.

14. Yao F, Yang S, Wang Z, Wang X, Ye J, Wang X, et al. Microbial taxa distribution is associated with ecological trophic cascades along an elevation gradient. Front Microbiol 2017; 8.

15. Na X, Xu TT, Li M, Ma F, Kardol P. Bacterial diversity in the rhizosphere of two phylogenetically closely related plant species across environmental gradients. J Soils Sediments 2017; 17: 122–132.

16. Jangid K, Williams MA, Franzluebbers AJ, Schmidt TM, Coleman DC, Whitman WB. Land-use history has a stronger impact on soil microbial community composition than aboveground vegetation and soil properties. Soil Biol Biochem 2011; 43: 2184–2193.

17. Talbot JM, Bruns TD, Taylor JW, Smith DP, Branco S, Glassman SI, et al. Endemism and functional convergence across the North American soil mycobiome. Proc Natl Acad Sci 2014; 111: 6341–6346.

18. Oono R, Rasmussen A, Lefèvre E. Distance decay relationships in foliar fungal endophytes are driven by rare taxa: distance decay in fungal endophytes. Environ Microbiol 2017; 19: 2794–2805.

19. Amend AS, Cobian GM, Laruson AJ, Remple K, Tucker SJ, Poff KE, et al. Phytobiomes are compositionally nested from the ground up. PeerJ 2019; 7: e6609.

20. Fitzpatrick CR, Copeland J, Wang PW, Guttman DS, Kotanen PM, Johnson MTJ. Assembly and ecological function of the root microbiome across angiosperm plant species. Proc Natl Acad Sci 2018; 115: E1157–E1165.

21. Bálint M, Tiffin P, Hallström B, O’Hara RB, Olson MS, Fankhauser JD, et al. Host genotype shapes the foliar fungal microbiome of balsam poplar (*Populus balsamifera*). PLoS One 2013; 8: e53987.

22. Bálint M, Bartha L, O’Hara RB, Olson MS, Otte J, Pfenninger M, et al. Relocation, high-latitude warming and host genetic identity shape the foliar fungal microbiome of poplars. Mol Ecol 2015; 24: 235–248.

23. Coleman-Derr D, Desgarennes D, Fonseca-Garcia C, Gross S, Clingenpeel S, Woyke T, et al. Plant compartment and biogeography affect microbiome composition in cultivated and native *Agave* species. New Phytol 2016; 209: 798–811.

24. Lindow SE, Brandl MT. Microbiology of the phyllosphere. Appl Environ Microbiol 2003; 69: 1875–1883.

25. Redford AJ, Bowers RM, Knight R, Linhart Y, Fierer N. The ecology of the phyllosphere: geographic and phylogenetic variability in the distribution of bacteria on tree leaves. Environ Microbiol 2010; 12: 2885–2893.

26. Leff JW, Del Tredici P, Friedman WE, Fierer N. Spatial structuring of bacterial communities within individual *Ginkgo biloba* trees. Environ Microbiol 2015; 17: 2352–2361.

27. Thompson LR, Sanders JG, McDonald D, Amir A, Ladau J, Locey KJ, et al. A communal catalogue reveals Earth’s multiscale microbial diversity. Nature 2017; 551: 457–463.

28. de Souza RSC, Okura VK, Armanhi JSL, Jorrín B, Lozano N, da Silva MJ, et al. Unlocking the bacterial and fungal communities assemblages of sugarcane microbiome. Sci Rep 2016; 6: 28774.

29. Wearn JA, Sutton BC, Morley NJ, Gange AC. Species and organ specificity of fungal endophytes in herbaceous grassland plants. J Ecol 2012; 100: 1085–1092.

30. Lindström ES, Langenheder S. Local and regional factors influencing bacterial community assembly. Environ Microbiol Rep 2012; 2012: 1–9.

31. Kembel SW, O’Connor TK, Arnold HK, Hubbell SP, Wright SJ, Green JL. Relationships between phyllosphere bacterial communities and plant functional traits in a neotropical forest. Proc Natl Acad Sci 2014; 111: 13715–13720.

32. Frank AC, Saldierna Guzmán JP, Shay JE. Transmission of bacterial endophytes. Microorganisms 2017; 5: 70.

33. Beckers B, Op De Beeck M, Weyens N, Boerjan W, Vangronsveld J. Structural variability and niche differentiation in the rhizosphere and endosphere bacterial microbiome of field-grown poplar trees. Microbiome 2017; 5: 25.

34. Desgarennes D, Garrido E, Torres-Gomez MJ, Peña-Cabriales JJ, Partida-Martinez LP. Diazotrophic potential among bacterial communities associated with wild and cultivated *Agave* species. FEMS Microbiol Ecol 2014; 90: 844–857.

35. Fonseca-García C, Coleman-Derr D, Garrido E, Visel A, Tringe SG, Partida-Martínez LP. The cacti microbiome: interplay between habitat-filtering and host-specificity. Front Microbiol 2016; 7: 150.

36. Peay KG, Schubert MG, Nguyen NH, Bruns TD. Measuring ectomycorrhizal fungal dispersal: macroecological patterns driven by microscopic propagules. Mol Ecol 2012; 21: 4122–4136.

37. Wu Z, Raven PH, Hong D. Hibiscus tiliaceus. eFloras. 2019. Missouri Botanical Garden, St. Louis, MO & Harvard University Herbaria, Cambridge, MA, pp 287–288.

38. Motooka P, Castro L, Nelson D, Nagai G, Ching L. Weeds of Hawaiʻi’s pastures and natural areas: an identification and management guide. 2014. University of Hawaiʻi Press, Honolulu.

39. Quesada T, Hughes J, Smith K, Shin K, James P, Smith J. A low-cost spore trap allows collection and real-time PCR quantification of airborne *Fusarium circinatum* spores. Forests 2018; 9: 586.

40. Smith DP, Peay KG. Sequence depth, not PCR replication, improves ecological inference from next generation DNA sequencing. PLoS One 2014; 9: e90234.

41. Walters W, Hyde ER, Berg-Lyons D, Ackermann G, Humphrey G, Parada A, et al. Improved bacterial 16S rRNA gene (V4 and V4-5) and fungal internal transcribed spacer marker gene primers for microbial community surveys. mSystems 2016; 1: e00009–15.

42. Rivers AR, Weber KC, Gardner TG, Liu S, Armstrong SD. ITSxpress: Software to rapidly trim internally transcribed spacer sequences with quality scores for marker gene analysis. F1000Research 2018; 7: 1418.

43. Hannon GJ. FASTX-Toolkit: FASTQ/A short-reads pre-processing tools. 2010.

44. Rognes T, Flouri T, Nichols B, Quince C, Mahé F. VSEARCH: a versatile open source tool for metagenomics. PeerJ 2016; 4: e2584.

45. Callahan BJ, McMurdie PJ, Rosen MJ, Han AW, Johnson AJA, Holmes SP. DADA2: High resolution sample inference from Illumina amplicon data. Nat Methods 2016; 13: 581–583.

46. Callahan BJ, McMurdie PJ, Holmes SP. Exact sequence variants should replace operational taxonomic units in marker-gene data analysis. ISME J 2017; 11: 2639–2643.

47. Gdanetz K, Benucci GMN, Vande Pol N, Bonito G. CONSTAX: a tool for improved taxonomic resolution of environmental fungal ITS sequences. BMC Bioinformatics 2017; 18: 538.

48. Frøslev TG, Kjøller R, Bruun HH, Ejrnæs R, Brunbjerg AK, Pietroni C, et al. Algorithm for post-clustering curation of DNA amplicon data yields reliable biodiversity estimates. Nat Commun 2017; 8: 1188.

49. Schloss PD, Westcott SL, Ryabin T, Hall JR, Hartmann M, Hollister EB, et al. Introducing mothur: Open-Source, Platform-Independent, Community-Supported Software for Describing and Comparing Microbial Communities. Appl Environ Microbiol 2009; 75: 7537–7541.

50. Davis NM, Proctor DM, Holmes SP, Relman DA, Callahan BJ. Simple statistical identification and removal of contaminant sequences in marker-gene and metagenomics data. Microbiome 2018; 6: 226.

51. McMurdie PJ, Holmes S. Phyloseq: an R package for reproducible interactive analysis and graphics of microbiome census data. PLoS One 2013; 8: e61217.

52. Giambelluca TW, Shuai X, Barnes ML, Alliss RJ, Longman RJ, Miura T, et al. Evapotranspiration of Hawai‘i. 2014.

53. Hijmans RJ. Raster: geographic data analysis and modeling. 2019.

54. Anderson MJ. A new method for non-parametric multivariate analysis of variance. Austral Ecol 2001; 26: 32–46.

55. Oksanen J, Blanchet FG, Friendly M, Kindt R, Legendre P, McGlinn D, et al. Vegan: community ecology package. 2019.

56. Guillot G, Rousset F. Dismantling the Mantel tests. Methods Ecol Evol 2013; 4: 336–344.

57. Dormann CF, Gruber B, Fruend J. Introducing the bipartite package: analysing ecological networks. R News 2008; 8: 8–11.

58. Dormann CF, Fründ J, Blüthgen N, Gruber B. Indices, graphs and null models: analyzing bipartite ecological networks. Open Ecol J 2009; 2.

59. Atmar W, Patterson BD. The measure of order and disorder in the distribution of species in fragmented habitat. Oecologia 1993; 96: 373–382.

60. Almeida-Neto M, Guimarães P, Guimarães PR, Loyola RD, Ulrich W. A consistent metric for nestedness analysis in ecological systems: reconciling concept and measurement. Oikos 2008; 117: 1227–1239.

61. Zarraonaindia I, Owens SM, Weisenhorn P, West K, Hampton-Marcell J, Lax S, et al. The soil microbiome influences grapevine-associated microbiota. mBio 2015; 6.

62. Zinger L, Amaral-Zettler LA, Fuhrman JA, Horner-Devine MC, Huse SM, Welch DBM, et al. Global patterns of bacterial beta-diversity in seafloor and seawater ecosystems. PLoS One 2011; 6: e24570.

63. Fraser MW, Gleeson DB, Grierson PF, Laverock B, Kendrick GA. Metagenomic evidence of microbial community responsiveness to phosphorus and salinity gradients in seagrass sediments. Front Microbiol 2018; 9: 1703.

64. Massoni J, Bortfeld-Miller M, Jardillier L, Salazar G, Sunagawa S, Vorholt JA. Consistent host and organ occupancy of phyllosphere bacteria in a community of wild herbaceous plant species. ISME J 2020; 14: 245–258.

65. Treseder KK, Maltz MR, Hawkins BA, Fierer N, Stajich JE, McGuire KL. Evolutionary histories of soil fungi are reflected in their large-scale biogeography. Ecol Lett 2014; 17: 1086–1093.

66. Nguyen HDT, Chabot D, Hirooka Y, Roberson RW, Seifert KA. *Basidioascus undulatus*: genome, origins, and sexuality. IMA Fungus 2015; 6: 215–231.

67. Becraft ED, Woyke T, Jarett J, Ivanova N, Godoy-Vitorino F, Poulton N, et al. Rokubacteria: genomic giants among the uncultured bacterial phyla. Front Microbiol 2017; 8.

68. Meiser A, Bálint M, Schmitt I. Meta-analysis of deep-sequenced fungal communities indicates limited taxon sharing between studies and the presence of biogeographic patterns. New Phytol 2014; 201: 623–635.

69. Crowther TW, Maynard DS, Crowther TR, Peccia J, Smith JR, Bradford MA. Untangling the fungal niche: the trait-based approach. Front Microbiol 2014; 5: 579.

70. Shakya M, Gottel N, Castro H, Yang ZK, Gunter L, Labbé J, et al. A multifactor analysis of fungal and bacterial community structure in the root microbiome of mature *Populus deltoides* trees. PLoS One 2013; 8: e76382.

