## Supplemental Information for "Multiscale interactions between plant part and a steep environmental gradient determine plant microbial composition in a tropical watershed"

#### Supplemental methods

*Detailed sampling methods*

Aside from the air microhabitat type, we collected all samples with sterile cotton swabs (Puritan® Diagnostics LLC, Guilford, ME, USA). At each of the nine *Hibiscus tiliaceus* trees along our transect, the adaxial and abaxial surfaces of two leaves were sampled. Four different petioles, independent from the leaves sampled, were swabbed between the stem and the leaf base. Two axils of the stem and main branches were swabbed for 10 seconds on each individual. Near the main trunk of each tree, two stems with a circumference of approximately 3 cm were selected for sampling. The entire surface area of the selected stem was swabbed in 10 cm long sections. Root (rhizosphere) samples were collected by selecting an area of the root with diameter 1.2–3.8 cm, and buried 1–8 cm below the surface. The roots were unearthed carefully to not break the epidermis; then a length of 15–18 cm was swabbed across the root surface. The entire abaxial and adaxial surfaces were swabbed of one leaf found in the leaf litter under the sampled individual at each site. Leaf litter samples were standardized across sites by selecting *H. tiliaceus* leaves which met the following criteria: leaves found directly underneath branches of the *H. tiliaceus* tree, leaf size roughly 6.35 × 6.35 cm, and leaf decomposition no more than 10% of total surface area. Once per site, we used a tubular soil sampler (1.5 cm diameter) to dig a 5-cm hole 1 m from extent of the canopy, which we used as an estimate to avoid sampling the rhizosphere. If the *H. tiliaceus* was adjacent to a stream, the sampling hole was on the opposite side of the tree to avoid runoff. We swabbed the sides and bottom of the hole. To avoid contamination, we sterilized the soil sampler between sites with 70% EtOH.

Air samplers were assembled following Quesada *et al.* (2018) with some modifications. The assembly procedure of one air sampler is as follows: A loop was made at the middle by twisting a 1.3 mm (16-gauge) thickness straightened metal rod about 20–25 cm long. Two standard microscopic slides (Thermo Fisher Scientific, Waltham, MA, USA) were secured at two edges of the metal rod by using all-purpose glue and cable ties. Microscopic slides were surface sterilized with 10% bleach followed by 70% ethanol solutions inside a laminar flow. Then the slides were coated with a thin layer of silicone vacuum grease with a cotton swab. We then cut microtiter plate sealing film (Thermo Fisher Scientific, Waltham, MA, USA) into 5.0 × 1.5 cm, which we positioned on the grease while exposing the bottom (white removable) layer of the film. Prepared traps were carefully placed in sterilized plastic bags until deployment in the field. At the site, each trap was assembled using battery-powered rotating garden motor (In the Breeze®, Bend, OR, USA). A pair of sterilized forceps was used to peel off the removable layer from the film so that the adhesive side remained exposed. The trap carefully hung on a branch using a metal rod approximately 1.2–1.5 m from the ground. The motor was covered with a plastic bag and an aluminum dish to protect from rain, wind and debris. The setup was kept in the field for 2 weeks prior to sample collection. During sample collection we used sterile forceps to remove the film from the slides and place in a 2 mL twist cap microcentrifuge tube, pre-filled with 1 mL of lysis buffer.

*PCR Parameters*

PCR parameters for ITS samples were as follows: 95°C for 3 min for denaturation; 35 cycles of 95°C for 20 sec, 53°C for 15 sec, 72°C for 30 sec; and 72°C for 3 min for final elongation. PCR parameters for 16S were as follows: 95°C for 3 min for denaturation; 35 cycles of 95°C for 20 sec, 50°C for 15 sec, 72°C for 30 sec; and 72°C for 3 min for final elongation.

### References

Quesada T, Hughes J, Smith K, Shin K, James P, Smith J. A low-cost spore trap allows collection and real-time PCR quantification of airborne *Fusarium circinatum* spores. *Forests* 2018; **9**: 586.

*Supplemental figures*

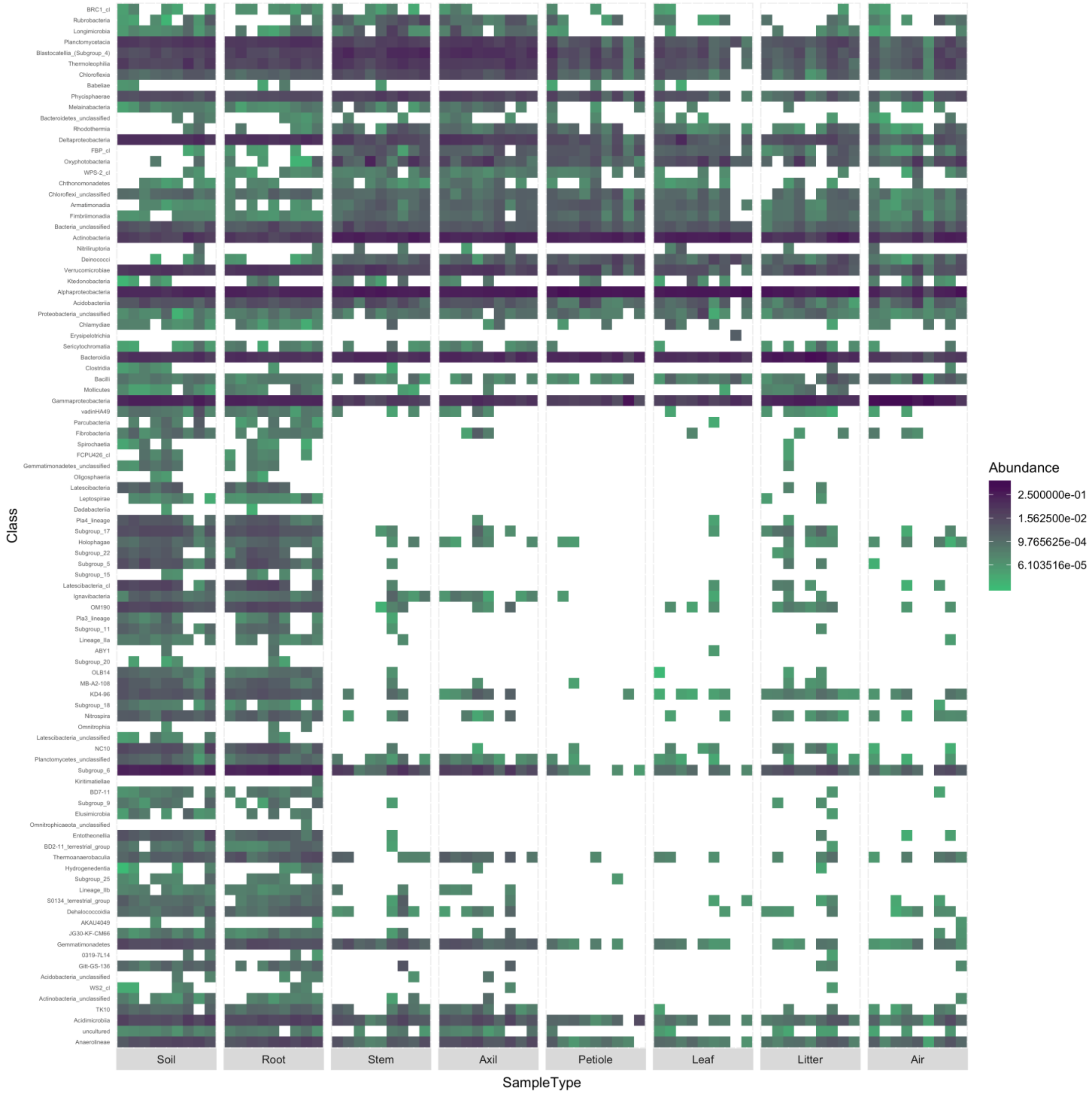


**Supplemental Figure S1:** Heatmap of bacterial classes inhabiting *Hibiscus tiliaceus* in Waimea Valley, O‘ahu. Colors indicate abundance. Within each tissue microhabitat (i.e., sample type), there are 9 blocks that represent sites along the environmental gradient, the leftmost being lowest in elevation and the rightmost being highest.

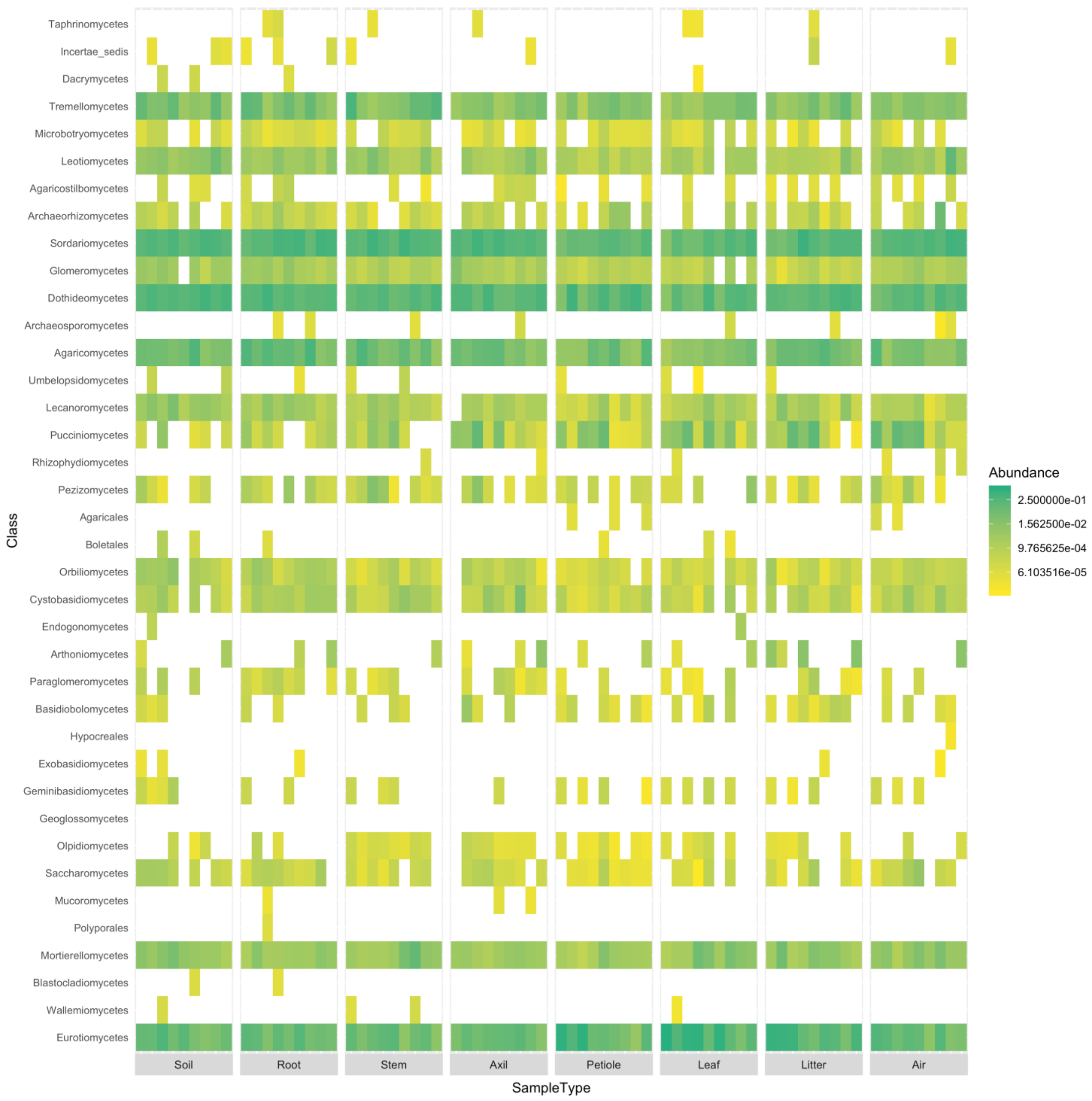


**Supplemental Figure S2:** Heatmap of fungal classes inhabiting *Hibiscus tiliaceus* in Waimea Valley, O‘ahu. Colors indicate abundance. Within each tissue microhabitat (i.e., sample type), there are 9 blocks that represent sites along the environmental gradient, the leftmost being lowest in elevation and the rightmost being highest.

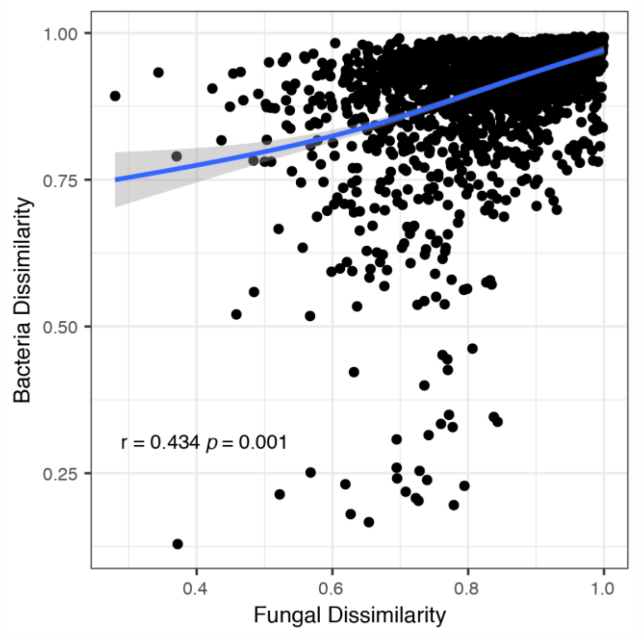


**Supplemental Figure S3:** Mantel test of Bray-Curtis dissimilarity and geographic distance between bacterial and fungal communities inhabiting *Hibiscus tiliaceus* in Waimea Valley, O‘ahu.

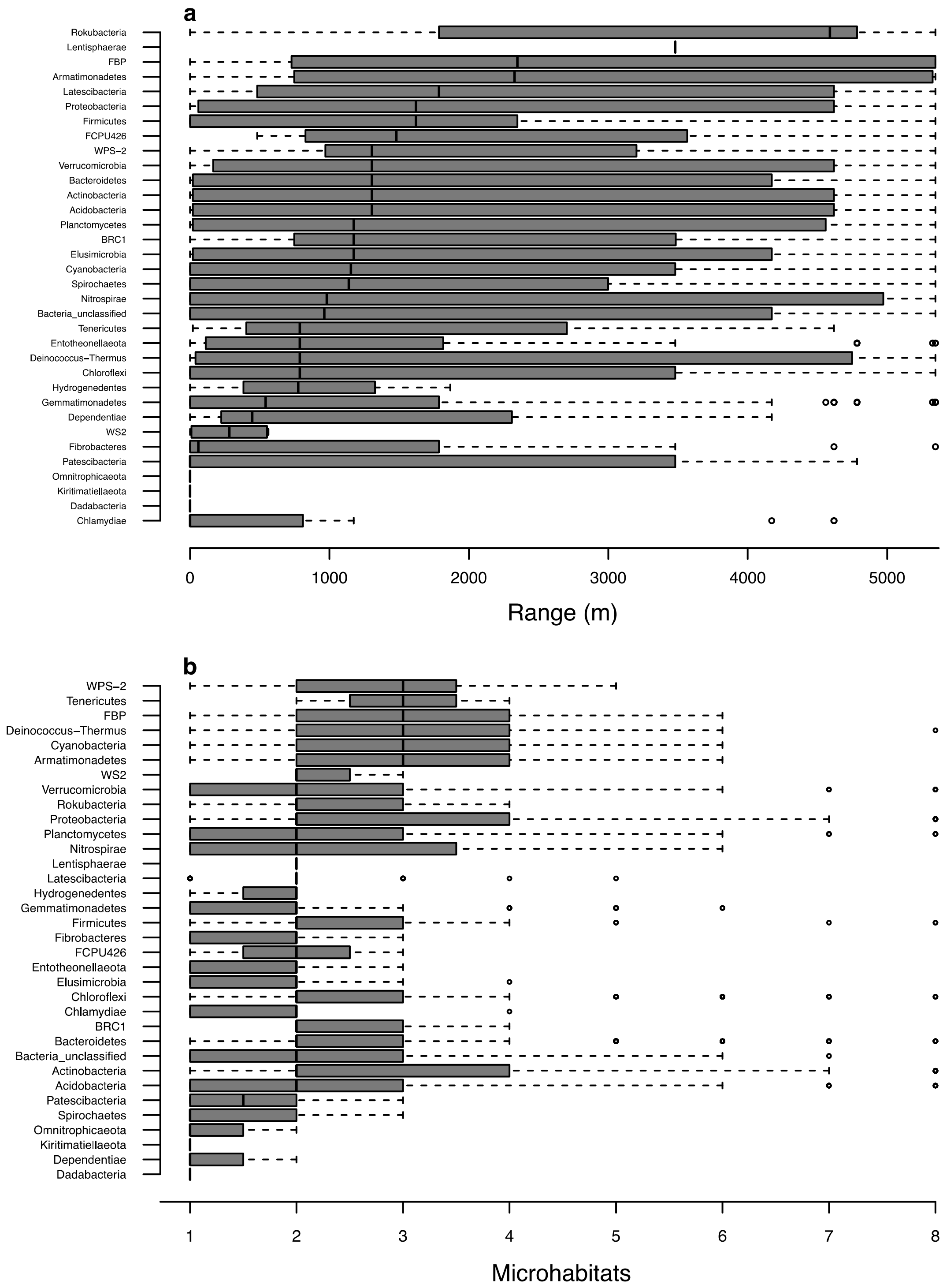


**Supplemental Figure S4:** Spatial distribution of bacterial phyla on *Hibiscus tiliaceus* across Waimea Valley, Oʻahu. (**a**) The range span (m) of each phylum across the sampling gradient (9 sites), and (**b**) the number of microhabitats where each bacterial phylum was observed.

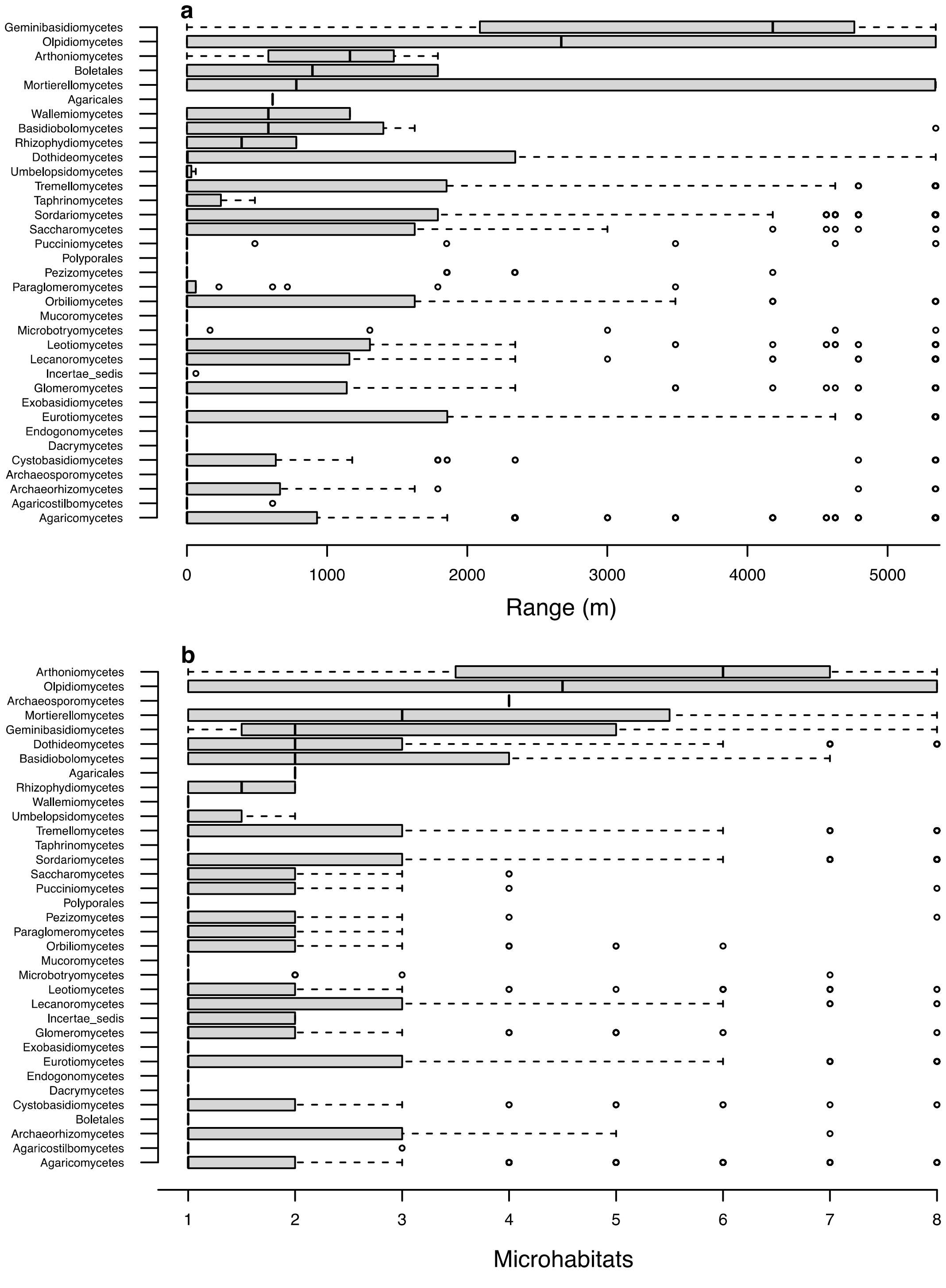


**Supplemental Figure S5:** Spatial distribution of fungal classes on *Hibiscus tiliaceus* across Waimea Valley, O‘ahu. (**a**) The range span (m) of each class across the sampling gradient (9 sites), and (**b**) the number of microhabitats where each fungal class was observed.
